# *Bradyrhizobium diazoefficiens* USDA110 nodulation of *Aeschynomene afraspera* is associated with atypical terminal bacteroid differentiation and suboptimal symbiotic efficiency

**DOI:** 10.1101/2020.11.24.397182

**Authors:** Quentin Nicoud, Florian Lamouche, Anaïs Chaumeret, Thierry Balliau, Romain Le Bars, Mickaël Bourge, Fabienne Pierre, Florence Guérard, Erika Sallet, Solenn Tuffigo, Olivier Pierre, Yves Dessaux, Françoise Gilard, Bertrand Gakière, Istvan Nagy, Attila Kereszt, Michel Zivy, Peter Mergaert, Benjamin Gourion, Benoit Alunni

## Abstract

Legume plants can form root organs called nodules where they house intracellular symbiotic rhizobium bacteria. Within nodule cells, rhizobia differentiate into bacteroids, which fix nitrogen for the benefit of the plant. Depending on the combination of host plants and rhizobial strains, the output of rhizobium-legume interactions is varying from non-fixing associations to symbioses that are highly beneficial for the plant. *Bradyrhizobium diazoefficiens* USDA110 was isolated as a soybean symbiont but it can also establish a functional symbiotic interaction with *Aeschynomene afraspera*. In contrast to soybean, *A. afraspera* triggers terminal bacteroid differentiation, a process involving bacterial cell elongation, polyploidy and membrane permeability leading to loss of bacterial viability while plants increase their symbiotic benefit. A combination of plant metabolomics, bacterial proteomics and transcriptomics along with cytological analyses was used to study the physiology of USDA110 bacteroids in these two host plants. We show that USDA110 establish a poorly efficient symbiosis with *A. afraspera*, despite the full activation of the bacterial symbiotic program. We found molecular signatures of high level of stress in *A. afraspera* bacteroids whereas those of terminal bacteroid differentiation were only partially activated. Finally, we show that in *A. afraspera*, USDA110 bacteroids undergo an atypical terminal differentiation hallmarked by the disconnection of the canonical features of this process. This study pinpoints how a rhizobium strain can adapt its physiology to a new host and cope with terminal differentiation when it did not co-evolve with such a host.

**Importance:** Legume-rhizobium symbiosis is a major ecological process in the nitrogen cycle, responsible for the main input of fixed nitrogen in the biosphere. The efficiency of this symbiosis relies on the coevolution of the partners. Some legume plants, but not all, optimize their return-on-investment in the symbiosis by imposing on their microsymbionts a terminal differentiation program that increases their symbiotic efficiency but imposes a high level of stress and drastically reduce their viability. We combined multi-omics with physiological analyses to show that the non-natural symbiotic couple formed by *Bradyrhizobium diazoefficiens* USDA110 and *Aeschynomene afraspera* is functional but displays a low symbiotic efficiency associated to a disconnection of terminal bacteroid differentiation features.

## Introduction

Nitrogen availability is a major limitation for plant development in many environments, including agricultural settings. To overcome this problem and thrive on substrates presenting a low nitrogen content, crops are heavily fertilized, causing important environmental damage and financial drawbacks^1,2^. Plants of the legume family acquired the capacity to form symbiotic associations with soil bacteria, the rhizobia, which fix atmospheric nitrogen for the plants’ benefit. These symbiotic associations lead to the development of rhizobia-housing root organs called nodules. In these nodules, the rhizobia adopt an intracellular lifestyle and differentiate into bacteroids that convert atmospheric dinitrogen into ammonia and transfer it to the plant. Critical recognition steps occur all along the symbiotic process and define the compatibility of the plant and bacterial partners^3^. While the mechanisms involved at the early stages of the symbiosis are well described, those of the later stages are much less clear and might affect not only the ability to interact but also the efficiency of the symbiosis (ie. the plant benefit).

Nodule-specific Cysteine-Rich (NCR) antimicrobial peptides produced by legumes of the Dalbergioids and Inverted Repeat Lacking Clade (IRLC) were proposed to play a crucial role in the control of host-symbiont specificity at the intracellular stage of the symbiosis^4^. NCR peptides are targeted to the bacteroids where they govern the bacteroid differentiation^5-9^. In these legumes, the differentiation process entails such profound changes that they suppress the bacteroids’ capacity to resume growth and is therefore referred to as terminal bacteroid differentiation (TBD). TBD contrasts with bacteroid formation in legumes that lack NCR genes (eg. soybean), where bacteroids are in a reversible state and can resume growth when released from nodules^10^. Specifically, TBD is associated with cell elongation, an increase in bacteroid DNA content through a cell cycle switch toward endoreduplication^6,9,11^. Furthermore, an increased permeability of the bacteroid envelope also occurs during TBD, most probably due to the interaction of NCR peptides with bacterial membranes ^6,7,10,12^. Together, these alterations of bacteroid physiology are associated to a strongly decreased viability of the differentiated bacteria, that fail to recover growth when extracted from nodules^6^.

While many rhizobia have a narrow host range, some species can nodulate a large array of plant species. One of them, *Bradyrhizobium diazoefficiens* USDA110, can trigger functional nodules without TBD on soybean (*Glycine max*), cowpea (*Vigna unguiculata*) or siratro (*Macroptilium atropurpureum*) (Fig 1A-B)^13^. In addition to these species, USDA110 induces also functional nodules on the TBD-inducing legume *Aeschynomene afraspera* (Fig. 1A-C)^14,15^. However, in *A. afraspera*, USDA110 shows only very limited features that are usually associated with TBD, suggesting that the bacterium might be resistant to the TBD process^16^.

**Figure 1.**
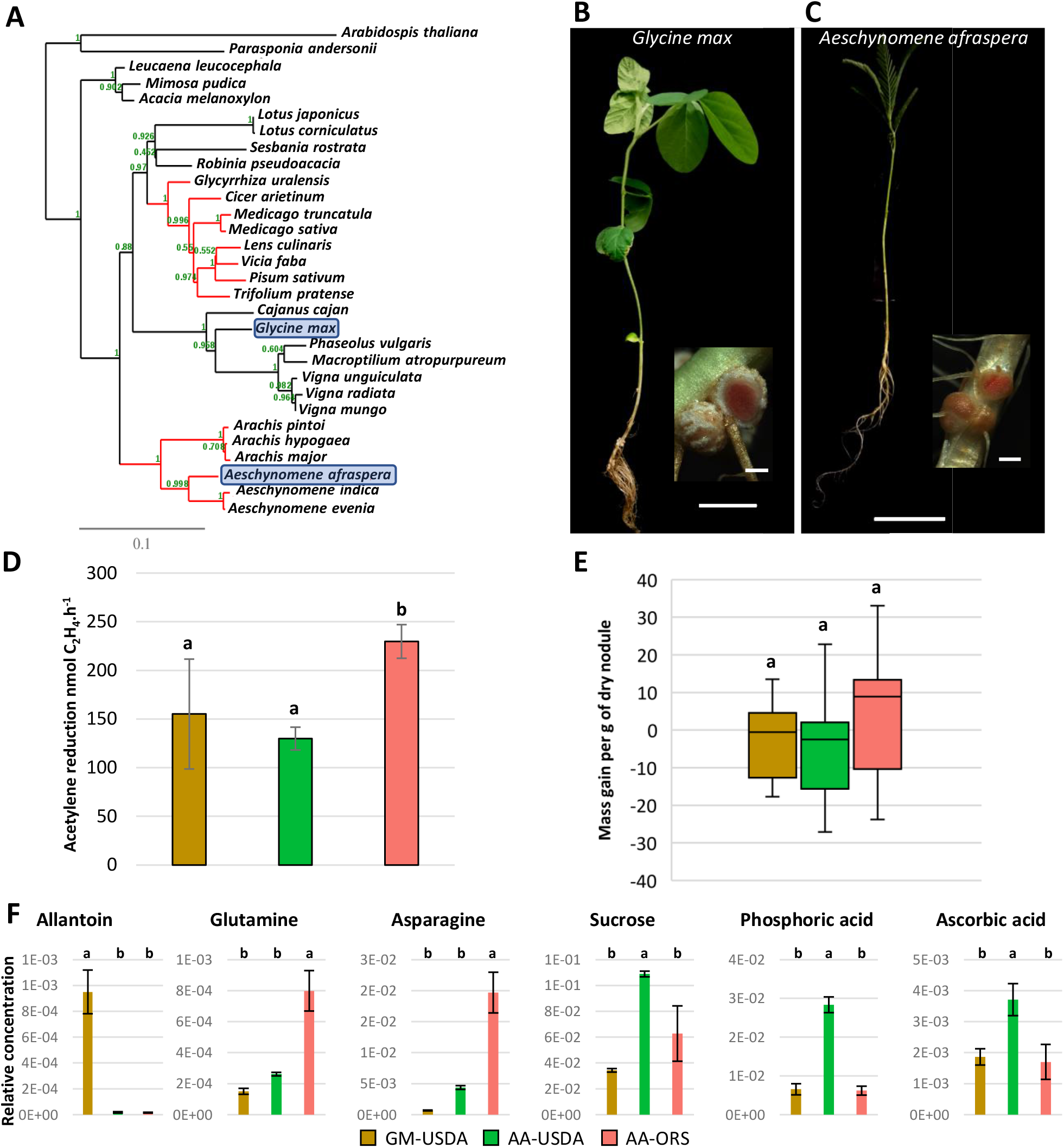
The non-adapted symbiotic couple formed by *Bradyrhizobium diazoefficiens* USDA110 and the NCR-producing plant *Aeschynomene afraspera* displays suboptimal nitrogen fixation and nodule metabolic dysfunction. A. Phylogenetic ML tree of a selection of plant species based on *matK* nucleotide sequences. Red branches indicate clades of legumes plants inducing terminal bacteroid differentiation. Blue boxes indicate the distantly-related host plants used in this study. Bootstrap values are mentioned in green on each node of the tree. B,C. General aspect of the plants and nodule sections (inlays) displaying the red coloration of leghaemoglobin of *G. max* (B) and *A. afraspera* (C) at 14 dpi. Scale bars: 5 cm (plants) and 1 mm (nodules). D, E. Nitrogen fixation activity determined by acetylene reduction assay (D) and gain in biomass attributable to the symbiosis (E) of 14 dpi plants. F. Whole-nodule metabolome determined by GC/MS or LC/MS at 14 dpi. Histograms show the average value of the relative metabolite concentration of four biological replicates. Letters represent significant differences after ANOVA and post hoc Tukey tests (p < 0.05). GM: *G. max* bacteroids, AA: *A. afraspera* bacteroids, USDA: *B. diazoefficiens* USDA110, ORS: *Bradyrhizobium sp*. ORS285.

Herein, we further characterized the bacteroid differentiation in the symbiosis between USDA110 and *A. afraspera*. Our observations, supported by whole-nodule metabolome analysis, indicate that USDA110 is poorly matched for nitrogen fixation with *A. afraspera*. To understand better the adaptation of *B. diazoefficiens* physiology to the *G. max* and *A. afraspera* nodule environment, we used a combination of transcriptomics (RNA-seq) and shotgun proteomics (LC-MS/MS) approaches. Finally, we find that USDA110 undergoes a terminal but atypical bacteroid differentiation in *A. afraspera* with a reduced cell viability and an increased membrane permeability, while cell size and ploidy level remain unchanged.

## Results

### *B. diazoefficiens* USDA110 is poorly matched with *A. afraspera* for nitrogen fixation

Previous reports indicate that *B. diazoefficiens* USDA110, the model symbiont of soybean, is able to establish a functional nitrogen-fixing symbiosis with *A. afraspera*, a phylogenetically distant host belonging to the Dalbergioid clade, which naturally interacts with photosynthetic rhizobia such as *Bradyrhizobium* sp. ORS285 (Fig. 1A-C)^14-18^. To evaluate the efficiency of this symbiosis, nitrogenase activity of USDA110- and ORS285-infected plants and their nitrogen content were determined. Although nitrogenase activity was detected in both types of nodules, it was significantly lower in USDA110-nodulated plants (Fig. 1D). A similar trend is observed for mass gain per nodule mass although this difference is not significant (Fig. 1E). Nitrogen and carbon contents seemedalso reduced in USDA110-nodulated plants as compared to ORS285-nodulated plants, reaching levels similar to those of non-inoculated plants (Fig. S1). Accordingly, ORS285-nodulated *A. afraspera* plants display darker green leaves than those interacting with USDA110.

Moreover, their shoot/root mass ratio, a metrics that reflects the nutritional status of the plant, is reduced in USDA110-nodulated *A. afraspera* plants as compared to ORS285-nodulated plants, indicating that the plant nutritional needs are not fulfilled (Fig. S2)^19^. To characterize further this suboptimal symbiosis, we analyzed the whole-nodule metabolome. Soybean nodules infected with USDA110 were used as a reference (Fig. S3). Allantoin, which is known to be the major nitrogen form exported from soybean nodules, is specifically detected in them (Fig. 1F)^20^. On the contrary, asparagine and glutamine are the principal exported nitrogen compounds in *A. afraspera* nodules and their amount is lower in USDA110-infected nodules as compared to ORS285-infected nodules, indicating a reduced nitrogen fixation by the bacteroids (Fig. 1F)^18^.

In addition, we find specifically in USDA110-infected *A. afraspera* nodules the accumulation of sucrose, phosphoric acid and ascorbate, and oppositely, a strong reduction in the trehalose content (Fig. 1F, Fig. S3). Sucrose derived from phloem sap is metabolized in nodules to fuel the bacteroids with carbon substrates, usually dicarboxylates. The accumulation of sucrose in nodules indicates a symbiotic dysfunction. Also, the accumulation of phosphoric acid in nodules suggests that nitrogen fixation is not reaching its optimal rate. Ascorbate has been shown to increase nitrogen fixation activity by modulating the redox status of leghemoglobin^21,22^. Thus, its accumulation in nodules with reduced nitrogen fixation capacity could be a stress response to rescue nitrogen fixation in nodules that do not fix nitrogen efficiently. A trehalose biosynthesis gene is upregulated in ORS285 bacteroids in *A. afraspera*, suggesting that TBD is accompanied by the synthesis of this osmo-protectant disaccharide^17^. The lower synthesis in USDA110 bacteroids suggests an altered TBD. Together these data indicate a metabolic disorder in the USDA110-infected nodules, in agreement with USDA110 being a suboptimal symbiont of *A. afraspera*.

### Overview of the USDA110 bacteroid proteomes and transcriptomes

In order to better understand the poor interaction between USDA110 and *A. afraspera*, the bacteroid physiology was analyzed through transcriptome and proteome analysis. The efficient soybean bacteroids and the free-living USDA110 cells cultivated in rich medium (exponential growth phase in aerobic condition) were used as references (Fig. 2A).

**Figure 2.**
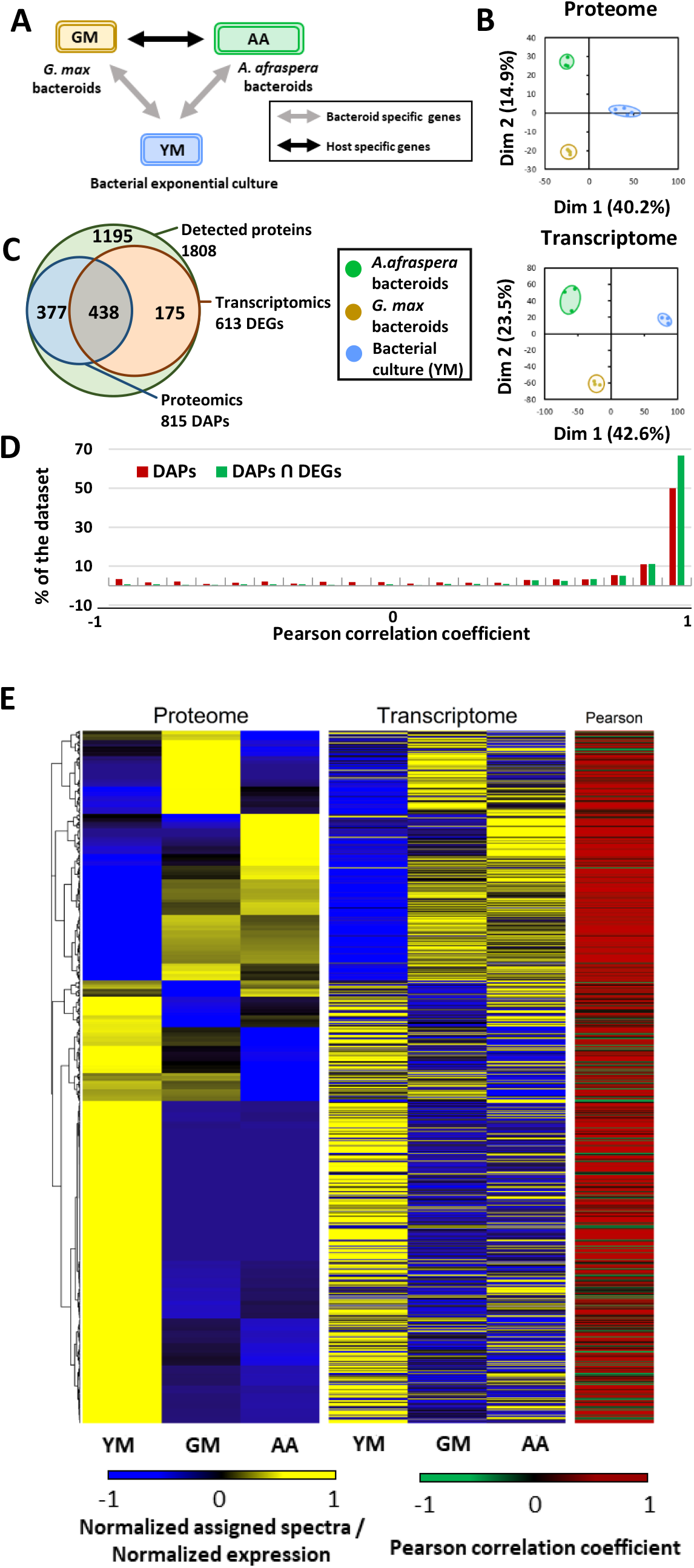
Experimental setup and general description of the trancriptomics and proteomics dataset. A. Experimental setup displaying the three biological conditions of this study. B. Principal component analysis of the proteomics and transcriptomics datasets. C. Venn diagram representing the overlap between differentially expressed genes (DEGs, FDR < 0.01 & |LFC| > 1.58) and differentially accumulated proteins (DAPs, FDR < 0.05) in at least one comparison and among the population of detected proteins. D. Pearson correlation coefficient (r) distribution between transcriptomic and proteomic datasets based on DAPs only (red) or DAPs that are also DEGs (green). E. Heatmaps and hierarchical clustering of the 815 DAPs and the corresponding transcriptomic expression values. The heatmaps show the standard score (z-score) of assigned spectra and DESeq2 normalized read counts, respectively. The color-coded scale bars for the normalized expression and value of Pearson correlation coefficient of the genes are indicated below the heatmap. YM: Yeast-Mannitol culture, GM: *G. max* bacteroids, AA: *A. afraspera* bacteroids.

Prior to quantification of transcript abundance or identification and quantification of protein accumulation, the transcriptome dataset was used to re-annotate the USDA110 genome with the EugenePP pipeline^23^. This allowed the definition of 876 new CDS, ranging from 92 to 1091bp (median size = 215bp or 71.6 aa) with 11.5% of them having a predicted function or at least a match using InterProScan (IPR). This extends the total number of CDS in the USDA110 genome to 9171. Moreover, we also identified 246 ncRNAs, ranging from 49 to 765 bp (median = 76 bp), which were not annotated before. Proteomic evidence could be found for 28 new CDS (3.2% of the new CDS, median size = 97.6 aa). The complete reannotation of the genome is described in Table S1.

In the proteome dataset, 1808 USDA110 proteins were identified. Principal component analysis (PCA) of all the replicate samples and sample types revealed their partitioning according to the tested conditions. The first axis of the PCA (40.2% of the observed variance) separates bacteroid profiles from the exponential culture, whereas the second axis separates *G. max* bacteroids from *A. afraspera* bacteroids (14.9% of the observed variance; Fig. 2B). The samples of the transcriptome datasets are similarly distributed on the PCA plot, with a first axis explaining 42.6% of the observed variance and a second axis explaining 23.5% of the observed variance (Fig. 2B).

Although differences are less pronounced in the proteome dataset than in the transcriptome dataset, COG analysis shows similar profiles across functional categories, except for membrane proteins that are less well identified in proteomics than transcriptomics (Fig. S4). In the transcriptomic dataset, 3150 genes are differentially expressed in at least one condition (differentially expressed genes or DEGs). Among the 1808 proteins identified, 815 show differential accumulation (differentially accumulated proteins or DAPs) and 438 of the cognate genes are also differentially expressed in the transcriptome datasets, whereas 175 DEGs are not DAPs (Fig. 2C).

We analyzed the Pearson correlation between transcriptomic and proteomic profiles and found that ∼66% of the bacterial functions that show significant differences in both approaches display a high correlation coefficient (r>0.9) whereas less than 1% of the functions show strong negative correlation (r<-0.9; Fig. 2D). This observation suggests that the transcriptome (which provides a more exhaustive view than the proteome) and the proteome provide a complementary picture of bacterial physiology, and they tend to show a congruent information if we restrict our analysis to the genes with differential accumulation/expression (Fig. 2E). However, there is still around 66% of the DEGs, which were detected by the proteomic analysis, that are not DAPs. Our description of the bacterial functions will be primarily based on the functions that are both DEGs and DAPs, as there is stronger evidence of their modulation in the tested conditions. The transcriptome alone will be used only when proteomics is not informative, for example to analyze regulons and stimulons that have been described previously in USDA110.

### Symbiotic functions common to both types of USDA110 bacteroids

Among the 815 DAPs, 705 and 699 proteins are significantly differentially accumulated in *G. max* and *A. afraspera* respectively compared to the bacterial culture control. Strikingly, 646 proteins are commonly differentially accumulated in both plant nodules (Table S1).

In the transcriptomic dataset, 1999 DEGs, representing ∼21% of the genome, were identified between the bacterial culture and the bacteroids, regardless of the host. Among them, 1076 genes displayed higher expression in nodules (including seven newly annotated ncRNAs and one newly annotated CDS among the 20 differentially expressed genes with highest fold change) and 923 genes were repressed *in planta* (including two newly annotated ncRNAs and two newly annotated CDS among the 20 DEG with highest fold change, Table S1).

Restricting the analysis to the bacterial functions that are both differentially expressed (DEG) and differentially accumulated (DAP) *in planta* in both hosts as compared to the bacterial culture identified 222 genes/proteins, 150 being upregulated and 72 being repressed *in planta* respectively (Fig. 3A). Notably, six newly annotated genes are in this gene list including one putative regulator (Bd110_01119) that is induced during symbiosis. Among the functions commonly DEG and DAP *in planta*, only four functions showed opposite trends in proteomics and transcriptomics.

**Figure 3:**
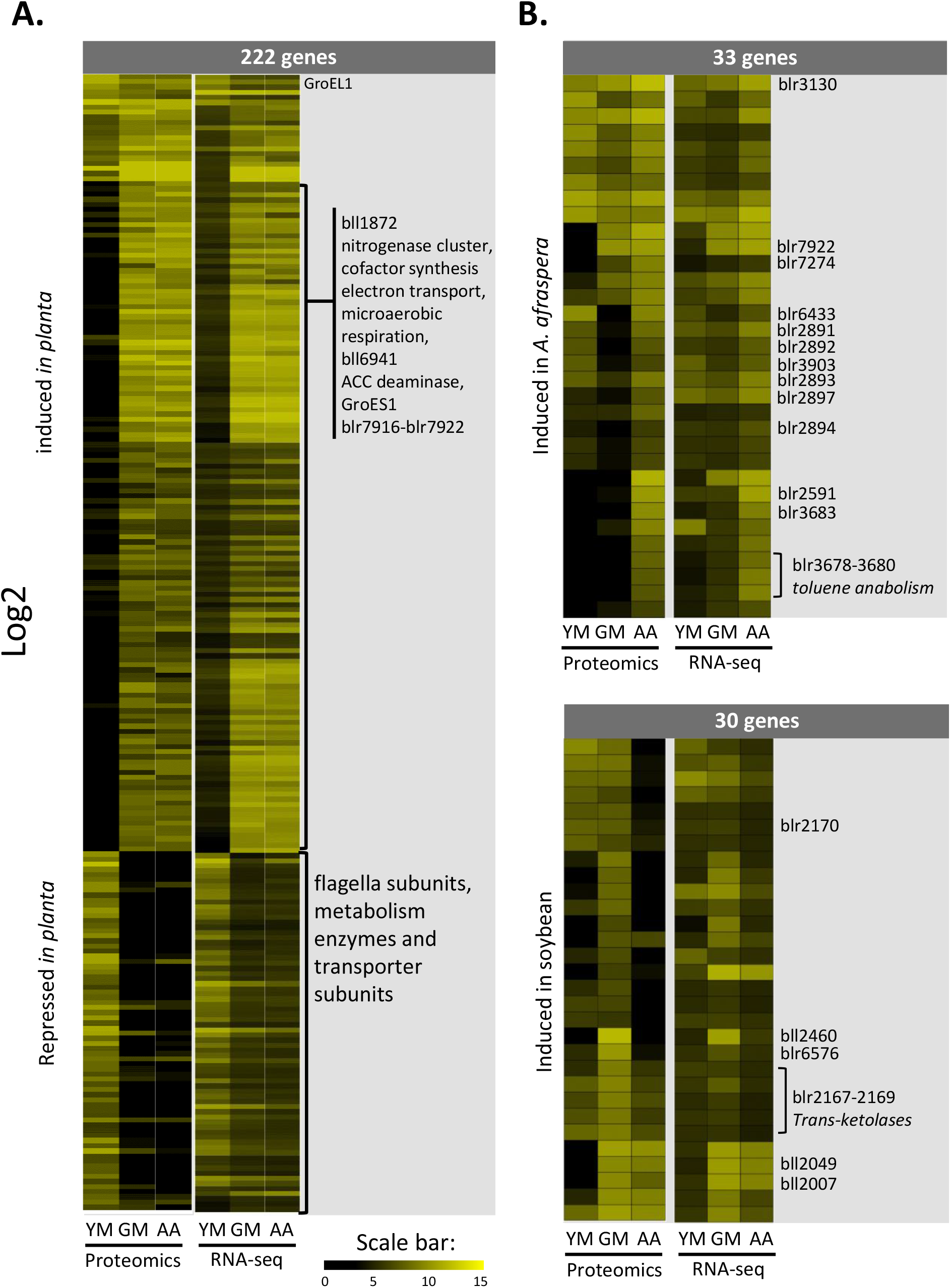
Symbiosis and host-specific functions that display congruency between transcriptomics and proteomics. A. Heatmap with SOM clustering displaying bacterial functions that are commonly DAP and DEG *in planta* in both host plants as compared to the culture reference. B. Heatmap displaying bacterial functions that are commonly DEG and DAP in one host as compared to the other (upper panel: *A. afraspera* > *G. max*; lower panel: *G. max* > *A. afraspera*). In panels A-B, data are presented as log 2 of DESeq2 normalized read counts (RNA-seq) or spectral counting (Proteomics). YM: Yeast-Mannitol culture, GM: *G. max* bacteroids, AA: *A. afraspera* bacteroids.

The proteome and transcriptome data provided a coherent view of the nitrogen fixation metabolism of *B. diazoefficiens* in the tested conditions. Key enzymes involved in microoxic respiration and nitrogen fixation were detected amongst the proteins having the highest spectra number in the nodule samples (Fig. 3A, Table S1) and the corresponding genes are among the most strongly expressed ones in bacteroids, while almost undetectable in the free-living condition. This includes for instance, the nitrogenase and the nitrogenase reductase subunits, which constitute the nitrogenase enzyme complex responsible for nitrogen conversion into ammonia. They belong to a locus of 21 genes from *blr1743* (*nifD*) to *bll1778* (*ahpC_2*), including the genes involved in nitrogenase cofactor biosynthesis, in electron transport to nitrogenase, and in microaerobic respiration, that are among the highest expressed ones in bacteroids of both host plants, both at the gene and protein expression level. The slightly higher level of the dinitrogenase reducatse NifH detected in proteomics was not supported by western blot analysis, which showed apparent similar protein level in both bacteroid conditions (Fig. S5). Strikingly, the two bacteroid types did not show a notable difference in the expression of these genes and proteins, suggesting that the activation of the nitrogen fixation machinery is not a limiting factor underlying the suboptimal efficiency of strain USDA110 in *A. afraspera* nodules.

In addition to these expected bacteroid functions, many other proteins were identified that specifically and strongly accumulated in both nodule types. This is the case of the chaperonins GroEL1/GroES1, which are strongly upregulated and reach high gene expression and protein levels in both bacteroids. The upregulation of these chaperonins is remarkable because other GroEL/GroES (4, 5 and 7) proteins are also very strongly accumulated in a constitutive manner. This indicates that bacteroids have a high demand for protein folding, possibly requiring specific GroEL isoforms, a situation reminding the requirement of one out of five GroEL isoforms for symbiosis in *Sinorhizobium meliloti*, the symbiont of *Medicago sativa*^12,24^. Another example of a bacteroid-specific function is the hydrogenase uptake system, whose gene expression was induced in both bacteroid types from nearly no expression in culture. Hydrogenase subunit HupL_2 (*bll6941*) was found amongst the proteins displaying the highest spectra number in the nodule samples suggesting important electron recycling in bacteroids of the two hosts. Another one is the 1-aminocyclopropane-1-carboxylic acid (ACC) deaminase (*blr0241*), which was also amongst the most strongly accumulated proteins in nodules and was significantly less abundant in free-living USDA110. An outer membrane protein (*bll1872*) belonging to the NifA regulon^25^ was also strongly induced *in planta*, with a transcript level among the top 10 genes in *A. afraspera*. Additionally, a locus of seven genes (*blr7916*-*blr7922*) encoding an amidase enzyme and a putative peptide transporter composed of two transmembrane domain proteins, two ATPases and two solute-binding proteins was strongly upregulated in the two bacteroid types, with three protein being also over-accumulated *in planta* (Fig. 3A; Table S1).

Oppositely, motility genes encoding flagella subunits (*bll5844*-*bll5846*), metabolic enzymes and transporter subunits are strongly downregulated during symbiosis and hardly detectable at the protein level *in planta* (Fig. 3A).

Taken together, these data show that both bacteroid types display a typical nitrogen fixation-oriented metabolism, with a partial shutdown of housekeeping functions. This indicates that despite the apparent reduced symbiotic efficiency of USDA110 in *A. afraspera* nodules, the bacterium fully expresses its symbiotic program within this non-native host as it does in soybean, its original host. Thus, the sub-optimal functioning of the *A. afraspera* nodules does not seem to come from a bacterial defect to express the symbiotic program, but possibly from an unfavorable host microenvironment or from a lack of metabolic integration of these maladapted partners.

### Host-specific functions

Comparison of the *A. afraspera* and *G. max* bacteroids revealed also significant differences in the proteomes and transcriptomes. At the transcriptomic level, 935 DEGs could be identified between the two bacteroid types (509 *A. afraspera* > *G. max* and 426 *G. max* > *A. afraspera*). One notable feature of the transcriptome is the identification of four newly annotated ncRNA and one new CDS amongst the 20 most induced DEGs in *A. afraspera* nodules and the presence of five newly annotated CDS amongst the 20 most induced DEGs in *G. max* nodules (Table S1). However, when considering only the functions that display congruent and significant differences in terms of transcripts and protein levels between plant hosts, we fall down to 63 genes/proteins, 33 being induced in *A. afraspera* nodules and 30 being induced in *G. max* nodules (Fig. 3B).

Interestingly, the phenylacetic acid degradation pathway (PaaABCDEIK, *blr2891-blr2897*) was highly expressed in *A. afraspera* nodules (although only PaaABCD and PaaK have been detected by proteomics), as well as a yet uncharacterized cluster of genes putatively involved in toluene degradation (*blr3675*-*blr3680*). The chaperone GroEL2 is also specifically induced in *A. afraspera*. Similarly, three S1 peptidases (Dop: *blr2591, blr3130* and *blr7274*) are highly expressed in the nodules of this latter host together with a RND efflux pump (*bll3903*) and a LTXXQ motif protein (*bll6433*), a motif also found in the periplasmic stress response CpxP^26^. The over-accumulation of these proteins suggests that bacteroids are facing stressful conditions during this interaction with *A. afraspera*. An uncharacterized ABC transporter solute binding protein (*blr7922*) was also overexpressed in *A. afraspera*.

One αβ hydrolase (*blr6576*) and a TonB-dependent receptor-like protein (*bll2460*) were over-accumulated in a *G. max*-specific manner. Similarly, an uncharacterized metabolic cluster including transketolases (*blr2167*-*blr2170*), the heme biosynthetic enzyme HemN1 (*bll2007*) and to a lesser extent an anthranilate phosphoribosyl-transferase (TrpD encoded by *bll2049*) are overexpressed in soybean nodules.

### USDA110 transcriptomics data in the perspective of previously described regulons and stimulons

USDA110 is one of the best-characterized rhizobial strains in terms of transcriptomic responses to various stimuli as well as the definition of regulons^27^. We analyzed the behavior of these previously defined gene networks in USDA110 in our dataset (Table S2). To initiate the molecular dialog that leads to nodule formation, plants secrete flavonoids like genistein in their root exudates, which are perceived by the rhizobia and trigger Nod factor production. At 14 dpi, when the nodule is formed and functioning, the genistein stimulon, which comprises the NodD1, NodVW, TtsI and LafR regulons, is not anymore activated in bacteroids. The symbiotic regulons controlled by NifA, FixK1, FixK2, FixLJ and sigma54 (RpoN) were activated *in planta*, indicating that nitrogen fixation was going on in both hosts. Accordingly, the nitrogen metabolism genes controlled by NtrC were activated *in planta*. Additionally, the PhyR/EcfG regulon involved in general stress response is not activated in bacteroids. Differences between hosts were however not observed for any of these regulons/stimulons. The only stimulon that showed differential expression between hosts is the one involved in aromatic compound degradation, which was highly expressed in *A. afraspera* nodules. Similar upregulation of the vanillate degradation pathway was observed in the transcriptome of *Bradyrhizobium* sp. ORS285 in *A. afraspera* and *A. indica* nodules^17^, suggesting that Dalbergioid hosts display a higher aromatic compound content in nodules than *G. max*. In line with this hypothesis, some of the most differentially accumulated sets of proteins (*A. afraspera* > *G. max*) are involved in the degradation of phenylacetic acid (PaaABCDK and *bll0339*) suggesting that the bacterium converts phenylalanine (or other aromatic compounds) ultimately to fumarate through this route (Fig. 3B)^28^. Similarly, enzymes of another pathway involved in phenolic compound degradation (*blr3675-blr3680*) are accumulated in *A. afraspera* nodules (Fig. 3B, Table S1).

### Expression pattern of orthologous genes between ORS285 and USDA110 in *A. afraspera* nodules

In a previous study^17^, a transcriptome analysis was performed on *Bradyrhizobium sp*. ORS285 in interaction with *A. afraspera* and in culture. *Bradyrhizobium* sp. ORS285 is a strain that co-evolved with *A. afraspera*, leading to an efficient symbiosis hallmarked by TBD, *id est* cell elongation and polyploidization of the bacteroids. In order to compare gene expression of these two nodule-forming rhizobia in culture and *in planta*, we determined the set of orthologous genes between the two strains using the Phyloprofile tool of Mage Microscope website. This analysis yielded a total of 3725 genes (Table S3). The heatmap on Figure 4A presents the modulation of gene expression (LFC) between *A. afraspera* nodules and the bacterial culture for the orthologous genes in each bacterium, regardless of their statistical significance. When taking FDR < 0.01 in account, we identified sets of genes that are differentially expressed *in planta* in either bacterium or in both (Fig. 4B).

**Figure 4:**
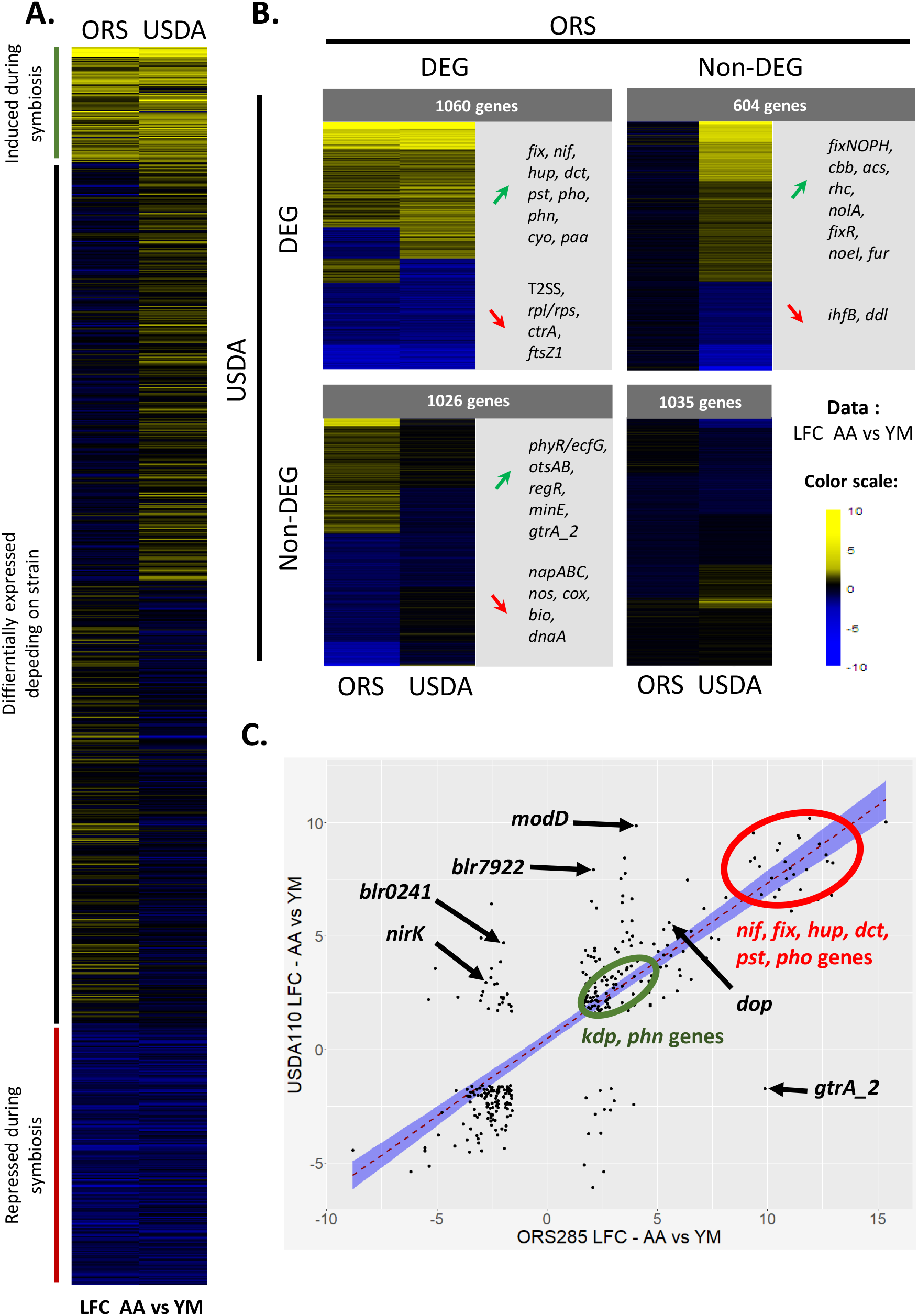
Expression pattern of *B. diazoefficicens* USDA110 and *Bradyhrizobium* sp. ORS285 orthologous genes *in planta* and in culture. A. Heatmap after SOM clustering of all the orthologous genes of USDA110 and ORS285 obtained with Phyloprofile. Values present the *in planta* LFC calculated after the read counts of the culture control versus *A. afraspera* 14 dpi nodules. B. Heatmaps of the orthologous genes after filtering on the FDR (< 0.01) values. Selected genes are highlighted for each class of interest. C. Dot plot of the orthologous genes that are DEG (FDR < 0.01 and |LFC| > 1.58) *in planta* (ie. in *A afraspera* nodules) in both strains. The red dashed line is for the linear regression and the blue envelope shows a 0.95 confidence interval of the linear regression.

Only 343 genes displayed differential expression (FDR < 0.01 and |LFC| > 1.58) *in planta* in both bacteria as compared to their respective culture control (Fig. 4C). A majority of these genes (86.8%) exhibited congruent expression patterns. First, the *nif, fix* and *hup* genes are commonly and highly induced in both strains during their symbiotic life with *A. afraspera*, a hallmark of a functional symbiosis. However, there are differences in their expression level, with a higher expression of the symbiotic genes in ORS285 (*nifHDK* represent 12.5% of all reads in *A. afraspera* nodules)^17^ than in USDA110 (*nifHDK* represent only 2.5% of all reads in *afraspera* nodules), consistently with a more efficient interaction occurring between ORS285 and *A. afraspera*. Additionally, the Kdp high affinity transport system, the phosphate (*pstCAB, phoU, phoE, phoC*) and phosphonate metabolism (*phnHIJKL*) are activated *in planta* in both bacteria (Fig. 4B-C). The stress-marker *dop* protease gene is also induced in both bacteria in *A. afraspera* nodules (Fig. 4C).

Additionally, 1026 genes were differentially expressed solely in ORS285, and similarly there was 604 DEG specific to USDA110 (Fig. 4B). For example, the general secretory pathway seems to be specifically induced in ORS285^17^. Oppositely, USDA110 displays an induction of the *rhcJQRU* genes which are involved in the injection of type three effector proteins that can be important for the establishment of the symbiosis whereas they are not induced or even repressed in ORS285 (Fig. 4B). This is also the case of the nitrite reductase encoding gene *nirK* (*blr7089*/BRAD285_v2_0763; Fig. 4C). In addition, USDA110 induces the expression of an ACC deaminase (*blr0241*), while its ortholog is repressed in ORS285 (BRAD285_v2_3570) during symbiosis (Fig. 4C). Bacterial ACC deaminases can degrade ACC, a precursor of ethylene, and thereby modulate ethylene levels in the plant host and promote the nodulation process^29^.

### *Bradyrhizobium diazoefficiens* USDA110 bacteroids undergo *bona fide* TBD in *Aeschynomene afraspera* nodules despite very weak morphological and ploidy modifications

In a previous description of the *A. afraspera* - *B. diazoefficiens* USDA110 interaction, the typical TBD features were not observed and the bacteroids were very similar to those in *G. max* where no TBD occurs^16^. At the molecular level, accumulation of the replication initiation factor DnaA is higher in soybean than in *A. afraspera* (Table S1). Similarly, the MurA peptidoglycan synthesis enzyme (encoded by *bll0822*) that may play a role in cell elongation during TBD was detected to similar levels in both bacteroids (Table S1). Taken together, the molecular data do not clearly indicate whether USDA110 bacteroids undergo TBD in *A. afraspera*. Therefore, we investigated the features of the USDA110 bacteroids in *A. afraspera* nodules in more detail.

We analyzed bacteroid differentiation features in USDA110 bacteroids extracted from soybean and *A. afraspera* nodules. The interaction between *A. afraspera* and *Bradyrhizobium* sp. ORS285 was used as a positive control for TBD features^9,30,31^. TBD is characterized by cell elongation. We quantified cell length, width, area and shape of purified bacteroids and culture controls. Whereas ORS285 bacteroids were enlarged within *A. afraspera* nodules as compared to their free-living conterparts, USDA110 bacteroids were similar to free-living bacteria in both soybean and *A. afraspera* (Fig. 5A; Fig. S6). Another feature of TBD is endoreduplication. Analysis of the bacterial DNA content of ORS285 bacteroids in *A. afraspera* by flow cytometry shows peaks at 6C and more^9^. As expected, USDA110 bacteroids in *G. max* yields only two peaks, at 1C and 2C, similarly to the cycling cells in the bacterial culture sample (Fig. 5B)^16^. Strikingly, similar results were obtained for USDA110 in *A. afraspera*. Thus, with respect to the DNA content and cell size, the USDA110 bacteroids do not display the typical TBD features in *A. afraspera* nodules. Loss of membrane integrity is a third hallmark of TBD that likely strongly contributes to the loss of viability of bacteroids. Time-course analysis of propidium iodide (PI) uptake by bacteroids and the corresponding culture controls were performed to assess bacteroid permeability (Fig. S7). Twenty minutes after PI application, USDA110 bacteroids from *A. afraspera* display an increased permeability that is much closer to ORS285 bacteroids in interaction with *A. afraspera* than to the low permeability of USDA110 bacteroids from *G. max* nodules (Fig. 5C). Also the free-living counterparts exhibit a very low permeability. Taken together, this suggests that the envelope of USDA110 bacteroids is more permeable in the NCR-producing *A. afraspera* nodules, even if it does not reach the permeability level of the ORS285 strain. To analyze bacterial viability, bacteroids extracted from nodules were plated and the colony forming units (cfu) were determined (Fig. 5D). In *G. max*, USDA110 formed 1.46×10^10^ colonies/mg nodule (∼100% survival). Oppositely, ORS285 formed only 5.42×10^7^ colonies/mg nodule in *A. afraspera* (∼0.5% survival). Interestingly, USDA110 formed 1.13×10^8^ colonies/mg nodule in *A. afraspera* (∼1% survival), indicating that, despite the absence of cell enlargment and endoreduplication USDA110 bacteroids lose their viability and undergo a *bona fide* terminal differentiation in *A. afraspera*. Thus, in the NCR-producing plant *A. afraspera*, USDA110 bacteroids display a disconnection of the four canonical TBD features (ie. cell size, ploidy level, membrane permeability and cell viability).

**Figure 5.**
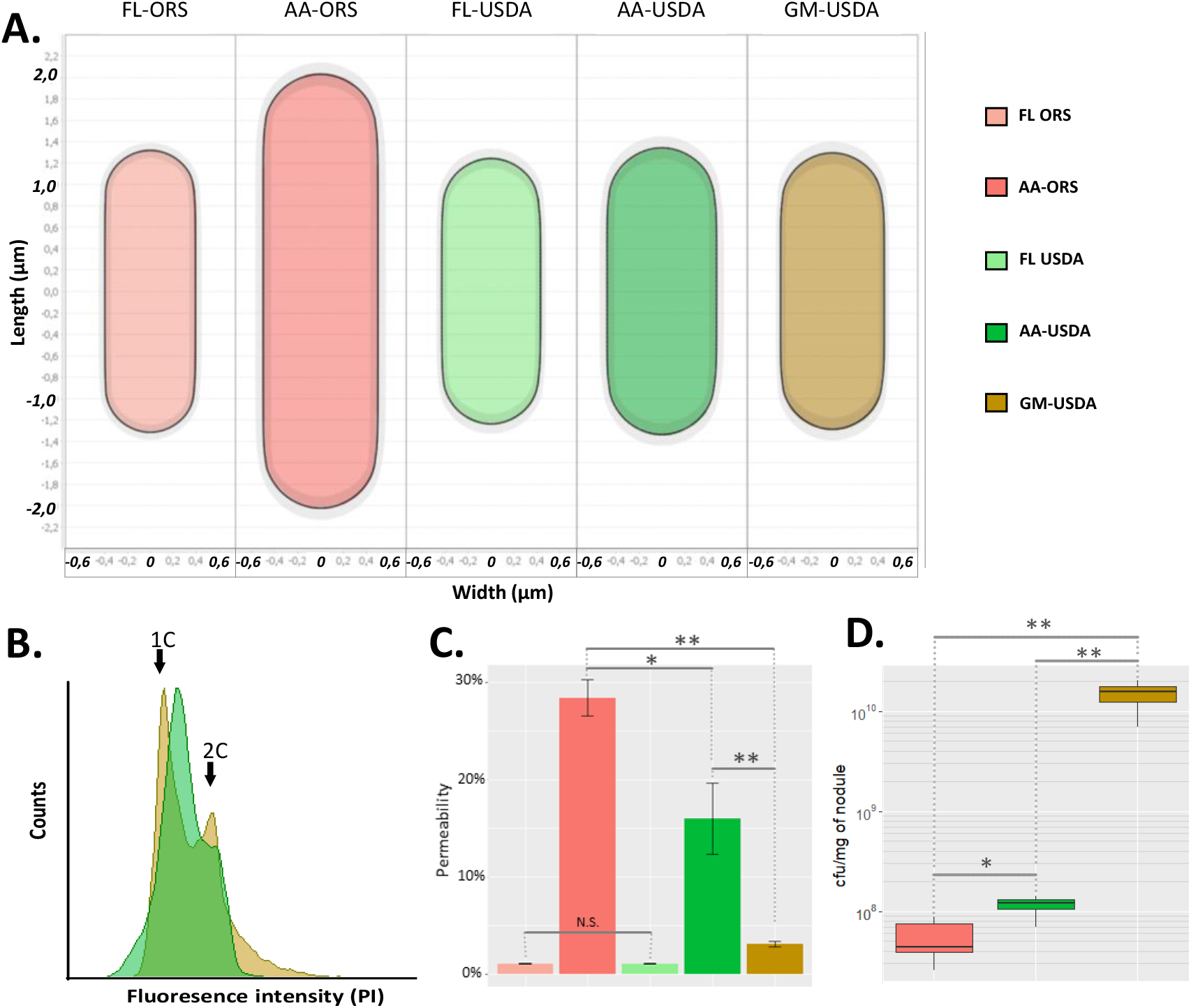
*B. diazoefficicens* USDA110 displays atypical bacteroid differentiation features in *A. afraspera* nodules. A. Average cell shape of free-living bacteria and bacteroids determined by MicrobeJ (900 < n < 21 000). B. DNA content of USDA110 bacteroids extracted from soybean and *A. afraspera* determined by flow cytometry. C. Assessment of the permeability of USDA110 and ORS285 free-living cells and bacteroids 20 min after PI treatment. *: wilcoxon test, p-value < 0.01. Five biological replicates were performed for bacteroids and two for free-living bacteria. D. Viability of soybean and *A. afraspera* extracted bacteroids at 14 dpi. Asterisks point out significant differences according to a wilcoxon test. *: p-value < 0.05; **: p-value < 0.01. Data are representative of 10 independent plants.

## Discussion

### *Aeschynomene afraspera* triggers atypical but terminal differentiation of USDA110 bacteroids

In a previous study, we noticed that, in *A. afraspera*, USDA110 forms a functional symbiosis although bacteroids do not display features that are usually associated with TBD^16^. Here we show that no endoreduplication and cell elongation of USDA110 occur in terminally differentiated bacteroids that fix nitrogen in a suboptimal way. Accordingly, the protein level of DnaA, the genome replication initiator, was higher in soybean than in *A. afraspera* bacteroids and the MurA level was not different between bacteroid conditions, confirming that polyploidization and cell elongation did not occur in this host. Such unusual terminal bacteroid differentiation is reminiscent of the bacteroids in *Glycyrrhiza uralensis*. This plant of the IRLC expresses NCR peptides^11^. However, one of its compatible symbionts, *Sinorhizobium fredii* strain HH103, does not undergo any loss of viability, no change in DNA content and no cell elongation^32^, while another symbiont, *Mesorhizobium tianshanense* strain HAMBI 3372 showed all TBD features^33^. The influence of the bacterial genotype on terminal/non-terminal differentiation of bacteroids was also suggested in *Medicago truncatula* in which, the gene *hrrP* might confer to some *Sinorhizobium* strains a resistance against the differentiation process triggered by some *M. truncatula* ecotypes^34^. In these two IRLC plants (ie. *M. truncatula* and *G. uralensis*), bacteria undergo a complete TBD or no TBD at all in a strain-dependent manner, but there is no clear uncoupling of the features of TBD (cell elongation/endoreduplication/altered viability) as shown here in the case of *B. diazoefficiens* USDA110-*A. afraspera*.

The surprising differentiation of USDA110 in *A. afraspera* nodules raises questions about the molecular mechanisms supporting this phenomenon. We consider two possible hypotheses: strain USDA110 might be more sensitive to the differentiation factors of the host than strain ORS285 and be rapidly “terminally” differentiated, before the other differentiation features, that are potentially important for symbiotic efficiency, can take place. Alternatively, USDA110 might be resistant to the plant effectors that trigger the elongation and polyploidization features.

In agreement with the latter possibility, the application of NCR peptides has very limited effect on strain USDA110 as compared to *S. meliloti* and to other plant-associated bacteria^16,35^. NCR insensitivity may be due to the thick hopanoid layer that is present in the outer membrane of strain USDA110, as the hopanoid biosynthesis mutant *hpnH* is more sensitive to NCR peptides and shows symbiotic defects in *A. afraspera* but not in *G. max*^36^. Moreover, the altered peptidoglycan structure in the strain USDA110 DD-carboxypeptidase mutant resulted in an increased TBD process with endoreduplicated and elongated bacteroids in *A. afraspera*^16^. This suggests that the envelope of strain USDA110 prevents a canonical TBD to occur. Possibly, NCR peptides are not able to reach their intracellular targets required to induce endoreduplication and cell division arrest, while their effect on cell viability through pore formation and membrane destabilization is still effective.

A survey of TBD in the legumes has identified multiple occurrences of the process in several subclades of the legumes but found that the majority of legumes do not have TBD^37^. The classification in this study was based on a morphological analysis of the bacteroids. Ancestral state reconstruction based on this classification suggested that the non-differentiated bacteroids are ancestral and that TBD evolved at least five times independently in legumes^37^. The discovery of bacteroids that are terminally differentiated without any obvious morphological changes opens the possibility that the occurrence of TBD might be underestimated in the legume family. Similarly, in the IRLC clade, the extent of morphological bacteroid differentiation was correlated to the size of the cationic NCR peptides repertoire and in legumes with few NCR peptides, the morphological modification of bacteroids can be minor^11,33^. In addition, at the molecular level, TBD is originally ascribed to the production of symbiotic antimicrobial peptides, the NCRs, by nodules^7^, but more recently, other types of antimicrobial peptides such as the NCR-like, GRP, MBP1 and CAPE peptides specifically produced in nodules of different plants were proposed to contribute to bacteroid differentiation^9,38-40^. Thus, if TBD would indeed be more widespread than currently estimated on the basis of morphological bacteroid features, the currently porposed evolutionary scenario of bacteroid formation might require revision.

### Terminal differentiation is associated with specific stress response

The TBD of strain USDA110 in *A. afraspera* is associated with a higher accumulation of stress markers compared to the *G. max* bacteroids. These markers include four proteases (Dop, Lon_2, *blr3130* and *blr7274*) and one chaperonin (GroEL_2). Similar induction of proteases and chaperonins have been reported in NCR-treated *S. meliloti* cultures^35^, indicating that this response may be linked to the perception of *A. afraspera* NCR-like peptides in USDA110.

The genes encoding these stress related proteins are not part of the well-characterized general stress response (GSR) controlled by the PhyR/EcfG signaling cascade in *B. diazoefficiens* USDA110^41^. On the other hand, we found that the PhyR/EcfG regulon in USDA110 is not activated in the bacteroids of both host plants (Table S2). This observation contrasts with our previous study of *Bradyrhizobium* sp. ORS285 transcriptome during symbiosis with *Aeschynomene* plants, which showed that the PhyR/EcfG cascade was upregulated *in planta*^17^. Nevertheless, the expression of the Dop protease was induced in *A. afraspera* in both bacteria (Fig. 4C). Together, the omics data suggest that bacteroids of *Bradyrhizobium* spp. activate stress-related genes in the TBD-inducing *A. afraspera* host but differences exist in the activation of specific stress responses at the strain level.

### Correlation between bacteroid differentiation features and symbiotic efficiency for the plant

TBD is associated with the massive production of symbiotic antimicrobial peptides such as NCR, NCR-like and CAPE peptides in different plants^5,9,38,40^. They represent ∼10% of the nodule transcriptomes in *M. truncatula* (analysis of the data from ref 42) and their production is thus potentially a strong energetic cost for the plant, raising questions about the benefits of the TBD process. TBD appeared independently in different legume clades^9,37^, suggesting that plants imposing this process obtain an advantage which might be a higher symbiotic benefit. Increased symbiotic efficiency has indeed been observed in hosts imposing TBD^17,43,44^. The findings reported here, comparing bacteroids and symbiotic efficiency in *A. afraspera* infected with strain ORS285 and strain USDA110, are in agreement with this hypothesis. Also in the symbiosis of *M. truncatula* in interaction with different *S. meliloti* strains, a similar correlation was observed between the level of bacteroid differentiation and the plant growth stimulation^45^. However, the simultaneous analysis of the bacteroid differentiation and symbiotic performance of an extended set of *Aeschynomene-Bradyrhizobium* interactions has shown that, perhaps not unexpectedly, the symbiotic efficiency of the plant-bacterium couple is not solely correlated with bacteroid differentiation and that other factors can interfere with the symbiotic efficiency as well^46^.

## Conclusion

*Bradyrhizobium diazoefficiens* USDA110 is a major model in the legume-rhizobium symbiosis, mainly thanks to its interaction with *G. max*, the worldwide most cultivated legume. Although omic studies have been conducted in this strain in symbiosis with various hosts^13,25^, this is the first time that this bacterium is studied at the molecular level in symbiosis with a NCR-producing plant that normally trigger a typical terminal bacteroid differentiation in its symbionts. The symbiosis between USDA110 and *A. afraspera* is functional even if nitrogen fixation and plant benefits are sub-optimal.

Terminal bacteroid differentiation is taking place in the NCR-producing host *A. afraspera*, as bacterial viability is impaired in USDA110 bacteroids, whereas morphological changes and the cell cycle switch to endoreduplication are not observed. We also show by combining proteomics and transcriptomics that the bacterial symbiotic program is expressed in *A. afraspera* nodules in a similar way as in *G. max*, although host-specific patterns were also identified. However, the bacterium is under stressful conditions in the *A. afraspera* host, possibly due to the production of NCR-like peptides in this plant. Integration of datasets from different bacteria in symbiosis with a single host, like ORS285 and USDA110 in symbiosis with *A. afraspera*, shed light on the differences in the stress responses activated in *A. afraspera* and confirmed that the symbiosis is functional but suboptimal in this interaction. The molecular data presented here provide a set of candidate functions that could be analyzed for their involvement in the adaptation to a new host and to the TBD process.

## Material and Methods

### Bacterial cultures and bacteroid extraction

*B. diazoefficiens* USDA110^47^ and *Bradyrhizobium* sp. ORS285 were cultivated in YM medium at 30°C in a rotary shaker^48^. For transcriptomic analysis, culture samples (OD_600nm_ = 0.5) were collected and treated as in Chapelle et al. (2015)^49^.

*G. max* ecotype Williams 82 and *A. afraspera* seeds were surface-sterilized and the plants were cultivated and infected with rhizobia for nodule formation as described in Barrière et al. (2017)^16^. Nodules were collected at 14 days post inoculation (dpi), immediately immersed in liquid nitrogen and stored at -80°C until use. Each tested condition (in culture and *in planta*) was produced in biological triplicates.

### Phylogeny analysis

Nucleotide sequence of *matK* genes were collected on NCBI using accession numbers described in references 50 and 51 and analyzed on phylogeny.fr (www.phylogeny.fr). They were aligned using ClustalW with manual corrections, before running a phyML (GTR - Gamma model) analysis with 500 bootstraps. A Bayesian inference tree was also generated (GTR + G + I) and provided similar topology as the maximum likelihood tree (data not shown). Trees were visualized and customized using TreeDyn.

### Genome annotation and RNA-seq analysis

Nodule and bacterial culture total RNA was extracted and treated as previously described in ^17^. Oriented (strand-specific) libraries were produced using the SOLiD Total RNA-seq kit (Life Technologies) and were sequenced on a SOLiD 3 station yielding ∼40 million 50bp single reads. Trimming and normalization of the reads were performed using the CLC workbench software. Subsequently, the reads were used to annotate the genome using EugenePP^23^, and the mapping was performed using this new genome annotation. Analysis of the transcriptome using DE-seq2 and data representation were performed as previously described^17^. Differentially expressed genes (DEG) showed an asbolute log_2_ fold change (|LFC|) > 1.58 (ie. fold change > 3) with a false discovery rate (FDR) < 0.01.

### Proteomic analysis

Bacteroids were extracted from 14 dpi frozen nodules^6^, while bacterial culture samples were collected as above, and the bacterial pellets were resuspended in -20°C acetone and lysed by sonication. Protein solubilization, dosage, digestion (trypsin 2% w/w) and solid phase extraction (using Phenomenex polymeric C18 column) were performed as described before^52^. Peptides from 800 ng of proteins were analyzed by LC-MS/MS with a Q Exactive mass spectrometer (Thermo Electron) coupled to a nanoLC Ultra 2D (Eksigent) using a nanoelectrospray interface (non-coated capillary probe, 10 µ i.d.; New Objective). Peptides were loaded on a Biosphere C18 trap-column (particle size: 5 μm, pore size: 12 nm, inner/outer diameters: 360/100 μm, length: 20 mm; NanoSeparations) and rinsed for 3 min at 7,5µl minute of 2% Acetonitrile (ACN), 0,1% Formic acid (FA) in water. Peptides were then separated on a Biosphere C18 column (particle size: 3 μm, pore size: 12 nm, inner/outer diameters: 360/75 μm, length: 300 mm; NanoSeparations) with a linear gradient from 5% of 0,1% FA in ACN (buffer B) and 95% of 0,1% FA in Water (buffer A) to 35% of buffer B and 65% of buffer A in 80 min at 300nl/min followed by a rinsing step at 95% of buffer B and 5% of buffer A for 6 min and a regeneration step with parameters of the start of the gradient for 8 min. peptide ions were analyzed using Xcalibur 2.1 software in data dependent mode with the following parameters: (I) full ms was acquire for the 400-1400 mz range at a resolution of 70000 with an AGC target of 3.10^6^; (ii) MS^2^ scan was acquired at a resolution of 17500 with an agc target of 5.10^4^, a maximum injection time of 120 ms and an isolation window of 3 m/z. The normalized collision energy was set to 27. MS^2^ scan was performed for the eight most intense ions in previous full MS scan with an intensity threshold of 1.10^3^ and a charge between 2 and 4. Dynamic exclusion was set to 50s. After conversion to mzXML format using msconvert (3.0.3706)^53^, data were searched using X!tandem (version 2015.04.01.1)^54^ against the USDA110 reannotated protein database and a homemade database containing current contaminants. In a first pass trypsin was set to strict mode and cysteine carbamidomethylation as a fixed modification and methionine oxidation, protein N-terminal acetylation with or without protein N-terminal methionine excision, N-terminal glutamine and carbamidomethylated cysteine deamidation, N-terminal glutamic dehydration as potential modifications. In a refine pass, Semi enzymatic peptides were allowed. Proteins inference was performed using X!TandemPipeline (version 3,4,3)^55^. A protein was validated with an E-value < 10^−5^ and 2 different peptides with an E-value < 0.05. Protein from contaminant database (*Glycine max* proteins and unpublished *Aeschynomene* Expressed Sequence Tags) were removed after inference. Proteins were quantified using the spectral counting method^56^. To discriminate differentially accumulated proteins (DAPs), ANOVA analysis was performed on the spectral counts and proteins were considered DAP when p-value < 0.05.

### Metabolomic analysis

Metabolites and cofactors were extracted from lyophilized nodules and analyzed by GC-MS and LC-MS respectively according to Su et al. (2016)^57^.

### Plant biomass and nitrogen fixation analysis

Dry mass of shoot, root and nodules was measured, and shoot-root mass ratio was calculated. The mass gain per g of dry nodule was calculated as the difference between total mean masses of the plants of interest and of the non-inoculated plants, divided by the mean mass of nodules. Thirty plants were used per condition. Nitrogenase activity was assessed by Acetylene Reduction Assay (ARA) on ten plants per condition as previously described^31^. The elemental analysis of leaf carbon and nitrogen content was performed as described in reference 18.

### Analysis of *B. diazoefficiens* USDA110 regulons and stimulons

Gene sets defined as regulons and stimulons were collected form the literature and the regulons/stimulons were considered as activated/repressed when ≥ 40% of the corresponding genes were DEG in a host plant as compared to the culture condition.

### Comparison of orthologous gene expression between *B. diazoefficiens* USDA110 and *Bradyhrizobium* sp. ORS285

The list of orthologous genes between USDA110 and ORS285 was determined using the Phyloprofile tool of the MicroScope-MAGE platform^58^, with identity threshold of 60%, maxLrap > 0 and minLrap > 0.8. The RNA-seq data from reference 17 and those of this study were used to produce heatmaps, for the genes displaying FDR < 0.01 (*A. afraspera* vs. YM), using R (v3.6.3) and drawn using pheatmap (v1.0.12) coupled with kohonen (v3.0.10) for gene clustering using the Self Organizing Maps (SOM) method. The DEG in both organisms (*A. afraspera* vs YM) were plotted for USDA110 and ORS285.

### Analysis of TBD features

Bacteroids were extracted from 14 dpi nodules and analyzed using a CytoFLEX S (Beckman-Coulter)^31^. For ploidy and live/dead analyses, samples were stained with propidium iodide (PI, ThermoFisher, 50 µg.mL^-1^ final) and Syto9 (ThermoFisher, 1.67 µM final). PI permeability was assessed over time on live bacteria. *Bradyrhizobium* sp. ORS285.pMG103-*nptII-GFP*^30^ and *B. diazoefficiens* USDA110 sYFP2-1^59^ strains were used to distinguish bacteroid from debris during flow cytometry analysis. For each time point, the suspension was diluted 50 times for measurement in the flow-cytometer. The percentage of bacteroids permeable to PI was estimated as the ratio of PI-positive over total bacteroids (GFP/YFP positive). Heat-killed bacteroids were used as positive control to identify the PI-stained bacteroid population.

For bacteroid viability assays, nodules were collected and surface sterilized (1 min NaClO 0.4%, 1 min 70% ethanol, two washes in sterile water). Bacteroids were subsequently prepared as previously described^31^ and serially diluted and plated (five µl per spot) in triplicate on YM medium containing 50 µg.mL^-1^ carbenicillin. Colony-forming units (cfu) were counted five days post plating and divided by the total nodule mass.

Bacterial cell shape, length and width were determined using confocal microscopy image analysis. Bacteroid extracts and stationary phase bacteria cultures we stained with 2.5 nM Syto9 for 10 minutes at 37°C and mounted between slide and coverslip. Bacteria imaging was performed on a SP8 laser scanning confocal microscope (Leica microsystems) equipped with hybrid detectors and a 63x oil immersion objective (Plan Apo, NA: 1.4, Leica). For each condition, multiple z-stacks (2.7µm width, 0.7 µm step) were automatically acquired (excitation: 488 nm; collection of fluorescence: 520-580 nm).

Prior to image processing, each stack was transformed as a maximum intensity projection using ImageJ software (https://imagej.nih.gov/ij/). Bacteria detection was performed with MicrobeJ (https://www.microbej.com/)^60^. First, bacteria were automatically detected on every image using an intensity based thresholding method with a combination of morphological filters (area: 1-20 µm^2^; length: 1 µm-∞; width: 0.5-1.3 µm) and every object was fitted with a “Rod-shaped” bacteria model. To ensure high data quality every image was manually checked to remove false positive (mainly plant residues) and include rejected objects (mainly fused bacteria). Then the morphology measurements and figures were directly extracted from MicrobeJ. ORS285 in culture and in symbiosis with *A. afraspera* were used as references for the analysis of TBD features.

### Western blot analysis

Detection of NifH by western blotting was performed using a commercial polyclonal antibody against a NifH peptide (Agrisera) respectively. The western blotting was carried out as previously described^61^ using bacterial exponential (OD_600nm_ = 0.5) and stationary (OD_600nm_ > 2.5) phase cultures as well as 14 dpi nodule-extracted bacteroids.

## Acknowledgments

The authors would like to thank Dora Latinovics for production and sequencing of the RNA-seq libraries and Mélisande Blein-Nicolas for her advices regarding the statistical analysis of the proteomic dataset. Q.N. and F.L. were supported by a PhD fellowship from the Université Paris-Sud. The present work has benefited from the core facilities of Imagerie-Gif (http://www.i2bc.paris-saclay.fr), member of IBiSA (http://www.ibisa.net), supported by ‘France-BioImaging’ (ANR-10-INBS-04–01), and the Labex ‘Saclay Plant Sciences’ (ANR-11-IDEX-0003-02). This work was funded by the Agence Nationale de la Recherche, grants n° ANR-13-BSV7-0013 and ANR-17-CE20-0011 and used resources from the National Office for Research, Development and Innovation of Hungary, grant n° 120120 to A.K.

## Authors’ contribution

QN, FL, PM, BGo and BA designed the work. QN, FL, AC, TB, MB, FGu, ES, ST and OP performed the experiments. QN, FL, MB, ES, YD, BGa, FGi, IN, AK, MZ, PM, BGo and BA analyzed the data. QN, FL, PM, BGo and BA wrote the paper.

**Supplementary figure and table legends**

**Figure S1. Nitrogen and carbon content in aerial parts of the plants were determined by elemental analysis**. GM: *G. max*, AA: *A. afraspera*, ORS: inoculated by *Bradyrhizobium* sp. ORS285, USDA: inoculated by *B. diazoefficiens* USDA110, NI: Non-inoculated plants.

**Figure S2. Nutritional status of 14 dpi plants determined by the shoot/root mass ratios**. AA: *A. afraspera*, ORS: inoculated by *Bradyrhizobium* sp. ORS285, USDA: inoculated by *B. diazoefficiens* USDA110, NI: Non-inoculated plants. Letters represent significant differences after t-test or ANOVA and post hoc Tukey tests (p < 0.05).

**Figure S3. Overview of the 129 quantified metabolites in *G. max* and *A. afraspera* whole nodules elicited by *B. diazoefficiens* USDA110 or *Bradyrhizobium* sp. ORS285**. Heatmap and hierarchical clustering of the 129 metabolites that were quantified either by gas-(GC-MS) or liquid-chromatography (LC-MS) coupled to mass spectrometry. Gm: *G. max*, Aa: *A. afraspera*, O: inoculated by *Bradyrhizobium* sp. ORS285, U: inoculated by *B. diazoefficiens* USDA110.

**Figure S4. General overview of the datasets using COG classification**. Repartitions of the assigned spectra (left panel) and normalized reads (right panel) among COG classes in the three conditions (blue: bacterial culture, ocher: *B. diazoefficiens* USDA110 in *G. max* nodules, green: *B. diazoefficiens* USDA110 in *A. afraspera* nodules).

**Figure S5. Western blot analysis of selected USDA110 proteins in culture and in bacteroids**. NifH protein were analyzed by western blots on purified USDA110 bacteroids extracted from soybean and *A. afraspera* nodules 14 dpi. Exponential and stationary phase cultures were used as controls.

**Figure S6. Analysis of cellular differentiation using automated morphometry**. A, B, C & D. Parameters were quantified by image analysis of syto9 stained bacteria and bacteroids using MicrobeJ. The process from raw images (A), segmentation (B), object detection (C) and measurements (D) is depicted with these four panels. E. Cell area. F. Cell width. G. Cell length.

**Figure S7. Kinetic analysis of bacterial membrane permeability**. Kinetics of propidium iodide uptake assays (reflecting membrane permeability) from which data presented in Figure 5C were extracted. The PI permeability was measured by flow cytometry over 60 min after treatment on *A. afraspera* nodule extracted USDA110 (AaU) or ORS285 (AaO) bacteroids and *G. max* extracted USDA110 bacteroids at 14 dpi (GmU). Exponential phase bacterial culture of USDA110 and ORS285 where used as controls. Each dot represent three independent measures and error bars represent the standard deviation of the samples.

**Table S1. Genome annotation, transcriptomic and proteomic data of *B. diazoefficiens* USDA110 generated in this study**. Description of proteomic and transcriptomic data of USDA110 related conditions. DESeq2 normalized reads, false discovery rate (FDR) values as well as log2 fold change (LFC) are used to describe transcriptomic data. On the other side, spectral counting (SC) along with related statistical indicators, Tukey statistical test result and p-value depict the proteomic data.

**Table S2. Expression analysis of selected *B. diazoefficiens* USDA110 regulons and stimulons**. Detailed analysis of the previously determined regulons and stimulons of USDA110 based on our transcriptomic data. A given regulon/stimulon was considered differentially regulated when ≥ 40% of the corresponding genes were differentially expressed in our conditions.

**Table S3. List of the 3725 orthologous genes shared by *B. diazoefficiens* USDA110 and *Bradyrhizobium* sp. ORS285 with their corresponding expression level in rich medium and in *A. afraspera* nodules**. This dataset was obtained after a Phyloprofile analysis on Mage Microscope website and was used to generate the Figure 4. Normalized read counts are shown together with the corresponding LFC and FDR as determined by DESeq2.

## Notes

### Competing Interest Statement

The authors have declared no competing interest.

### Summary of Updates

This manuscript was revised to include the supplemental tables that were not tranfered by the journal where the manuscript has been submitted.

