## Supplementary figures and images for "*Bradyrhizobium diazoefficiens* USDA110 nodulation of *Aeschynomene afraspera* is associated with atypical terminal bacteroid differentiation and suboptimal symbiotic efficiency"

### Supplemental Figures 1 to 7

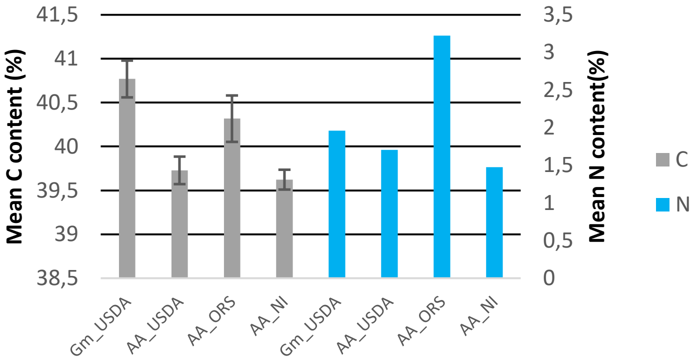

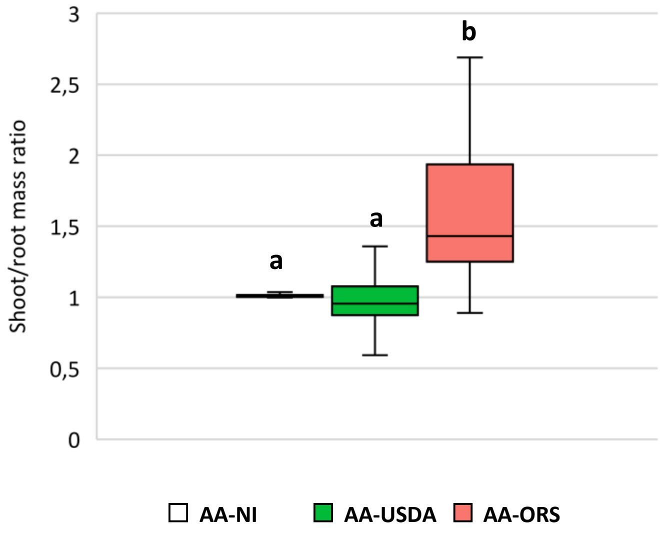


#### Figure S2

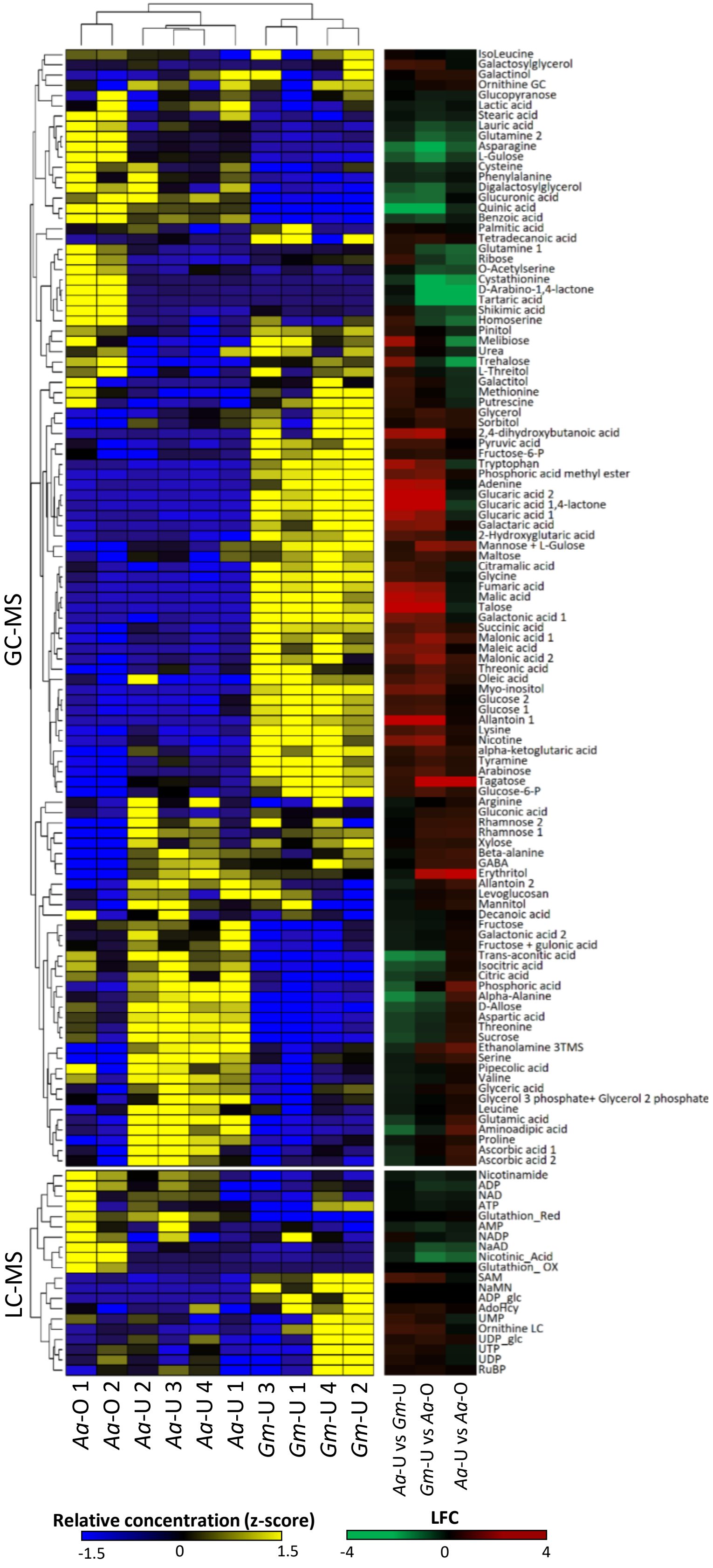

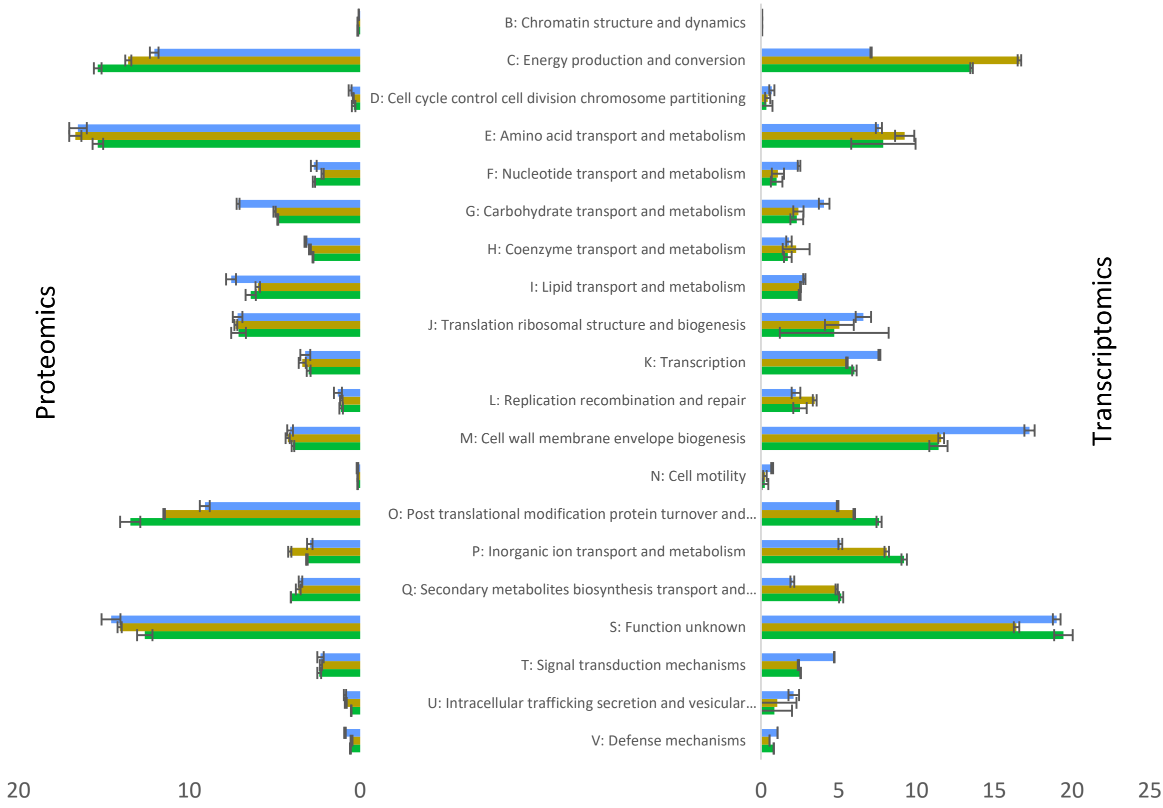

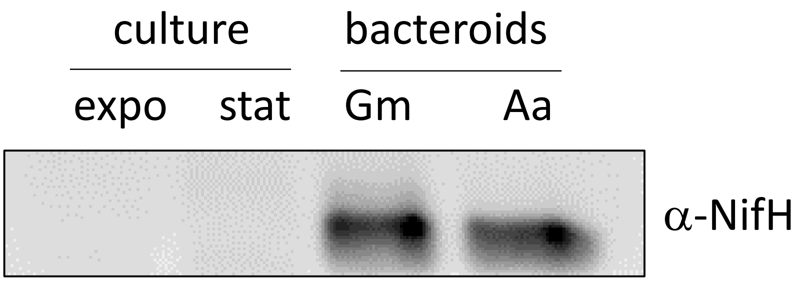


**Figure S5**

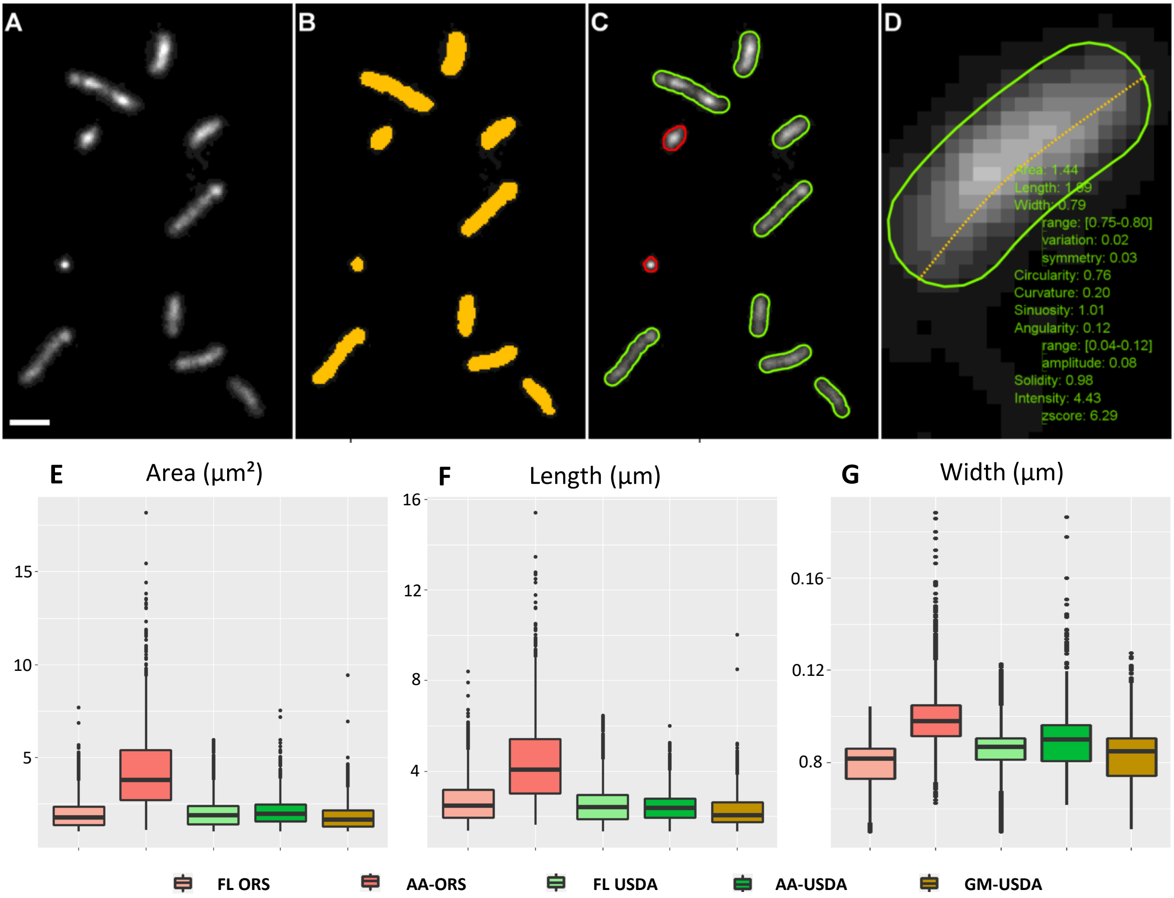

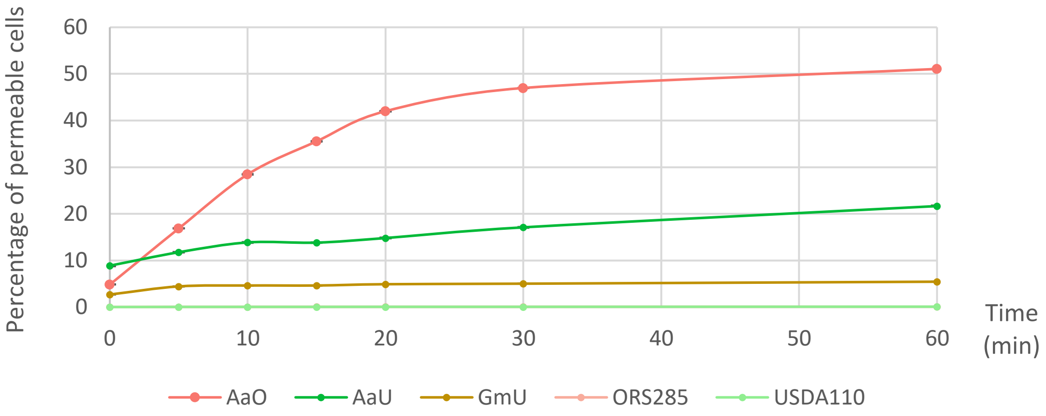
